# Flagellum malfunctions trigger metaboly as an escape strategy in *Euglena gracilis*

**DOI:** 10.1101/863282

**Authors:** Yong-Jun Chen

## Abstract

Euglenoids, a family of aquatic unicellular organisms, present the ability to alter the shape of their bodies, a process referred to as metaboly [1–5]. Metaboly is usually used by phagotrophic cells to engulf their prey. However, *Euglena gracilis* is osmotrophic and photosynthetic. Though metaboly was discovered centuries ago, it remains unclear why *E. gracilis* undergo metaboly and what causes them to deform [1–5], and some consider metaboly to be a functionless ancestral vestige [5]. Here, we show that flagellum malfunctions trigger metaboly and metaboly is an escape strategy adopted by *E. gracilis* when the proper rotation and beating of the flagellum are hindered by restrictions including surface obstruction, sticking, resistance, or limited space. Metaboly facilitates escape in five ways: 1) detaching the body from the surface and decreasing the attaching area attached to the interface, which decreases the adhering force and is advantageous for escaping; 2) enlarging the space between flagellum and the restricting surface which restores beating and rotation of the flagellum; 3) decreasing the torque of viscous resistance for rotation of the body and changing the direction of the body to restore flagellar function; 4) decreasing the length of the body, which pulls the flagellum away from the restrictive situations; and 5) crawling backwards on a surface or swimming backwards in a bulk fluid if the flagellum completely malfunctions or has broken off. Taken together, our findings suggest that metaboly plays a key role in enabling *E. gracilis* to escape from harmful conditions when flagellar functions is impaired or absent. Our findings can inspire the bionic design of adaptive soft robots and facilitate the control of water blooms of euglena in freshwater aquiculture industry.

---

*Euglena gracilis* is a species of flagellar unicellular protist [1–2]. The body of cells is covered by a pellicle [6], with a flagellum at the anterior tip. The regular rotation and beating of the flagellum pull the body in a spiral trajectory, and the cell body with a cigar shape rotates in a counterclockwise direction (as viewed from behind) [7,8]. By changing the mode of beating, the cell can change its direction and move in different trajectories in the free flagellum-driven motion [8]. Though *E. gracilis* is osmotrophic and photosynthetic traits, the cell retains metaboly of the body in the early evolution [3–8], which seems not crucial for the mobility. Metaboly is related to the cell pellicle, with strips on the pellicle surface controlling the plasticity of the cytoskeleton of the cell [1]. Though metaboly in *E. gracilis* was discovered by Harris in 1696 [3], the triggering mechanism and functions are still not clear. Recently, it is reported that metaboly in *E. gracilis* confined between two glass plates results in a crawling motion; however, it is unclear whether this function is relevant in the cells’ natural aquatic environment [5]. Therefore, why and under what conditions *E. gracilis* demonstrates metaboly remain open questions. The answers of these questions have implications for bionic designing of adaptive micro soft robots taking advantage of metaboly [9, 10] and water bloom control in freshwater aquiculture industry. Additionally, *E. gracilis*, at the bottom of the food chain, is the prey of the other cells, for eample, *Peraneama trichophorum* [11]. Escape strategy of *E. gracilis* was never investigated previously. Here, we observed *E. gracilis* motility under various ubiquitous conditions encountered in its natural living environments to determine the triggering mechanism and the role of metaboly when motility is restricted.

The air-water interface is the most important place for the cells to accept sunlight for photosynthesis, and is a preferred place for the cell in nature because of intensive sunlight due to phototaxis [12–14]. And various kinds of interface exist in aquatic environment. To determine when metaboly occurs, we observe *E. gracilis* behaviour under various conditions to mimic different types of restriction in its natural environment, including attaching under an air-water surface, attaching on a glass wall, moving in the vicinity of a contact line, moving in a viscous solution, bumping into an obstacle (another cell), and swimming in an extremely crowded environment. The cell showed metaboly when attaching under the air-water surface (Fig. 1a). When the cell tip contacted the air-water surface, the cells became trapped easily and attached under the surface (Fig. S1), which was never noted previously. Though the cells can glide on the surface by beating its flagellum on the free side of the cell (Fig. S2, see movies 6 and 7), we found that they could not detach from the air-water interface without metaboly. To escape from the surface, the cell deforms and rotates to change the direction, as shown in Fig. 1a. Whether the cell will undergo metaboly to try to detach from the surface depends on its intention. Similar behaviors were found when the cell was attached on a glass wall or a glass substrate. Fig. 1b shows the deformation and rotation of the cells induced by rotation and beating of the flagellum and escape from the attachment of a glass wall or substrate. Fig. 1c shows a process of metaboly and escape from a contact line mimicking boundaries of natural water. The cell tip contracts and rotates to turn the cell around. In the vicinity of a contact line (Fig. S3a), the air-water surface and the surface of the substrate forms a wedge-shaped space. The cell cannot change its direction easily and will exhibit metaboly. When there are many cells aggregating in the vicinity of the contact line, they will deform their bodies, rotate and try to escape away from the contact line (see movie 8). Whether it can swim away depends on the case. Sometimes, a small deformation to decrease its length is enough to turn the cell around (Fig. S3b). Usually, it will try metaboly and rotation several times. In a crowded environment, the cell deforms when it bumping into other individuals as shown in Fig. 1d. In particular, in extremely crowded conditions, all the cells show metaboly continuously (Fig. S4 and movie 9). Taken together, these findings suggest that metaboly occurs when the motion of the cells is hindered, whether by attachment; contact with a line, surface or obstacle; or resistance.

**Figure 1.**
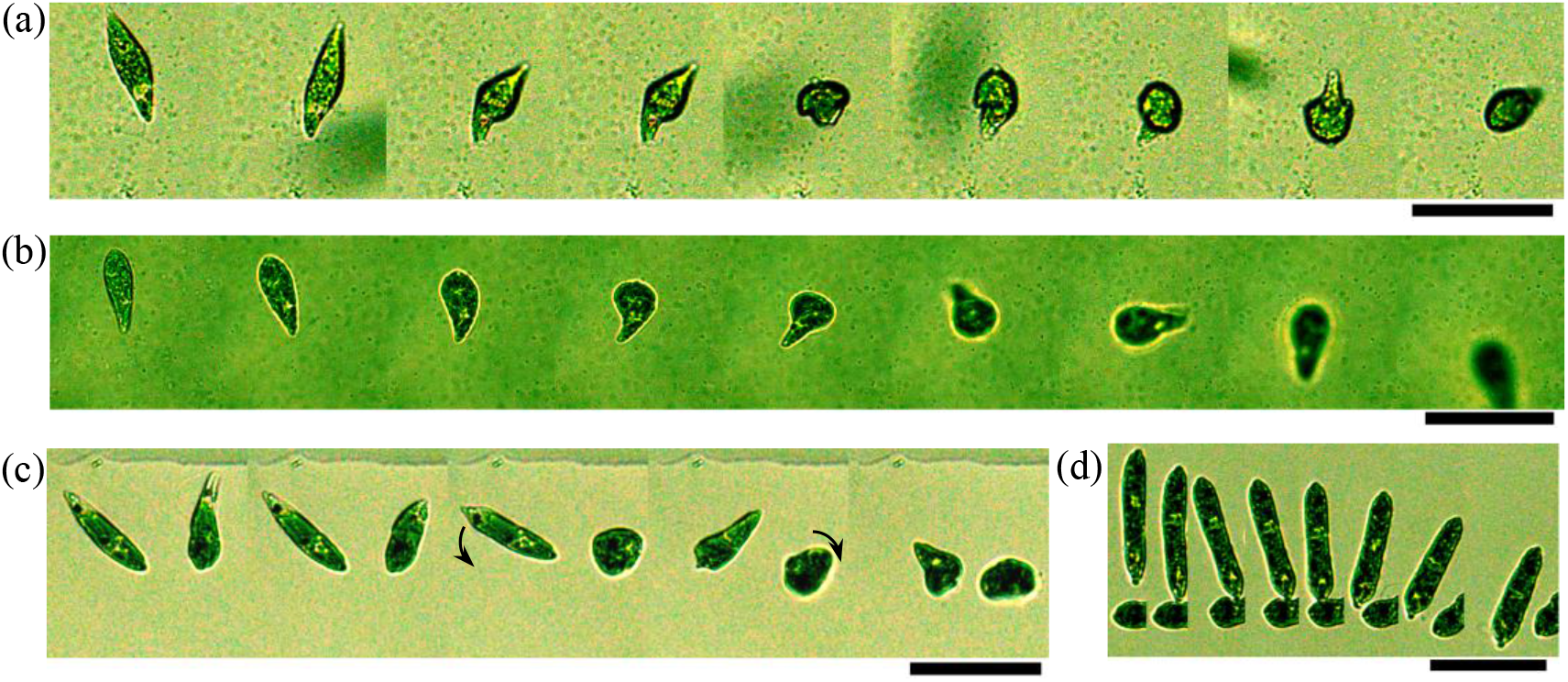
Metaboly and escape under various conditions. (a) At the air-water interface and escaping (movie 1). The air-water interface is at the top of the images. The time interval is 2.0 s. (b) Attached to a glass wall and escaping (movie 2). The glass wall is at the bottom of the images. The time interval is 2.0s. (c) Near the contact line and escaping (movie 3). The arrows indicate the directions of cell rotation. Time interval is 1.0 s. (d) Bumping into another cell and then decreasing the length of the body to escape (movie 4). Scale bars: 50μm.

To clarify what triggers *E. gracilis* metaboly, mechanical contact (with surface or other obstacles) on the pellicle or motion of flagellum, we observed the behaviours of the cells under highly viscous conditions (without contact with pellicle surface) by adding increasing amounts of sodium polyacrylate acid (PAAS) to the cell culture dispersion. We found *E. gracilis* survives well in PAAS solution. As shown in Fig. 2a, in the low viscosity (c<0.005 wt%) solution, the cells exhibited a cigar shape and moved fast, propelled by the flagella. At intermediate viscosity (0.0050 wt%<c<0.035 wt%), metaboly and flagellum-driven motion with a cigar shape coexist; and while some of the cells were propelled by their flagella, some exhibited metaboly. With increasing PAAS concentration in the range of 0.005 wt%<c<0.035 wt%, the fraction of cells demonstrating metaboly increases, the fraction of the flagellum-driven cells decreases, and the velocity of the flagellum-driven motion decreases. The exact value of the fraction of cells exhibiting metaboly is difficult to obtain because of limited visual field of microscope. In addition, one cell cannot always swim under flagellum-driven mode when the PAAS concentration is relatively high (0.035wt%≤c≤0.050wt%). No flagellum-driven motion was apparent when the concentration was larger than 0.050 wt%, and when the concentration was 0.10 wt %, the cells were immobilized by the highly viscous solution. Given the effect of resistance on the motion and shape of the cells, we next asked whether the increased viscosity affected not only the displacement velocity, but also the motion of the flagellum. Depending on the modes of flagellum motion, cells show various kinds of motion, including flagellum-driven displacement (the flagellum rotates and beats regularly, movie 12), rotation (the flagellum can rotate and beat, but not regularly, see Fig. 2b), swimming by metaboly with rotation (the abnormally beating flagellum cannot rotate, see Fig. 2b), and a mixture of flagellum-driven motion and metaboly (the flagellum intermittently failed to rotate and beat regularly, see movie 13). During rotation of the cell, the cell will deform and expand in the axis direction of the body repeatedly. During the metaboly swimming process, the trajectory of cell is not straight because of the rotation of the cell body. In highly viscous environments, the viscous resistance impedes the regular rotations and beating of the flagellum. The high viscosity of the solution not only decreases the velocity of the flagellum-driven motion, but increases the resistance for the rotation and beating of the flagellum. The coexistence of the flagellum-driven motion and metaboly suggests that metaboly is not triggered by the resistance force on the pellicle, as all the cells experience the same viscous drag on the surface during the motion. The metaboly in highly viscous condition and motion of cell corresponding to different modes of flagellum motion suggest that metaboly is related to the modes of flagellum motion while not mechanical contact on the pellicle.

**Figure 2.**
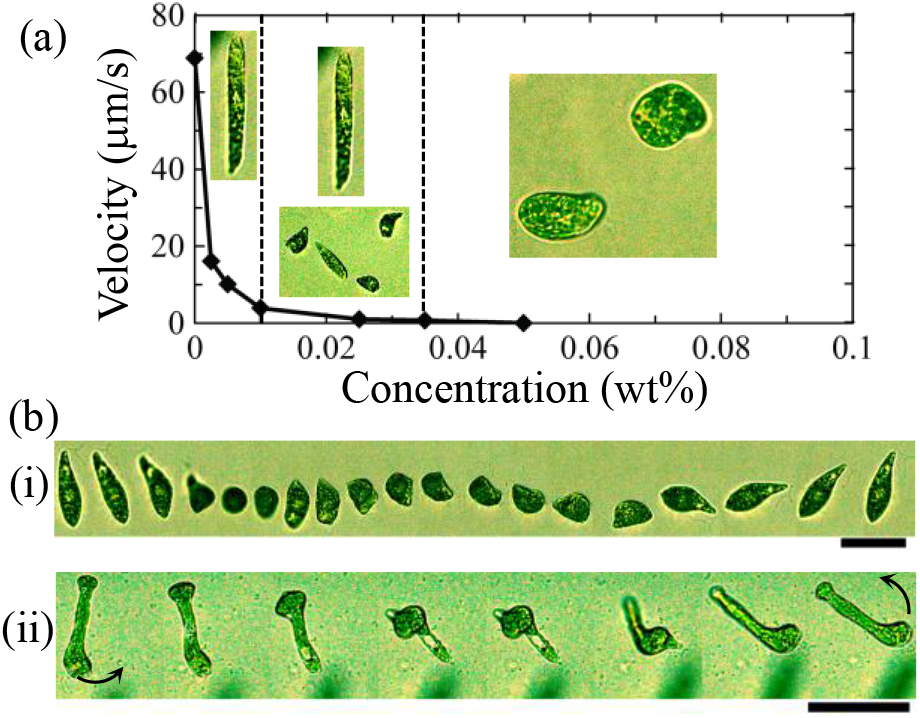
Transistion from flagellum-driven motion to metaboly in viscous solutions.(a) Conformation and velocity of *E. gracilis* at different concentrations of PAAS. Representative images of cell morphology are shown in the insets. The dashed lines indicate the regions of concentration. (b) E. gracilis mobility under viscous conditions: (i) Rotation and deformation of the cell (movie 10). Time interval is 0.5 s. (ii) Swimming by metaboly. Periodic deformation leads to swim and rotation when the flagellum cannot rotate but can beat (movie 11). The arrows indicate the directions of cell rotation. The time interval is 1.0 s. The concentration of the PAAS solution is 0.025wt%. Scale bars: 50μm.

Given that changes in the flagellar activity appeared to correlate with changes in cell shape, we next ask whether flagellar activity generally triggers metaboly in the ubiquitous aquatic conditions and metaboly affects flagellar function. To investigate this, we examine details in metaboly and flagellar motion. In highly viscous condition, the flagella of those cells exhibiting metaboly cannot beat and rotate regularly, while the flagella of those cells with a cigar shape are relatively stronger. We found that the flagellum was immobilized due to resistance, and that metaboly restored its ability to beat regularly (Fig. 3a). This process can also be clearly observed in the process of flagellum-driven motion with intermittent metaboly (movie 13). Another interesting finding is that the cells can rotate easily when the far end of the flagellum malfunctions and the flagellum cannot beat (Fig 3b). The rotation of the base end of the flagellum produces a torque on the cell body, and the cell changes it direction easily. During the rotation of the cell, metaboly is observed, and the cell has round shape (movie 15). When the cell nears or attached to a surface (including an air-water interface, a wall, or a substrate), the beating and rotation of flagellum will malfunction (Fig. 3c and d). When attached to a glass substrate, the flagellum can only beat on the left side (Fig. 3c, seen from above). When attached under the air-water surface, the flagellum can only beat on the right side of the cell body (same side with Fig. 3c, seen from above) (Fig. 3d). The reason is that the flagellum only rotates in the counterclockwise direction (when viewing the cell from behind), driven by a molecular motor. The air-water interface and substrate surface hinder the rotation and regular beating of the flagellum. The rotation of flagellum stops at the surface, and the beating can only continue in a two-dimensional space on one side of the cell (Fig. 3c and d). From the movies 6 and 7, we can see that the flow driven by the beating of the flagellum is unsymmetrical and that the fluid moves from the front to the left side. On the right side, the intensity of the flow is weak. Such unsymmetrical flow cannot produce a regular driving force to detach the cell body from the surface. The cell glides along the surface, and the body rotates within a two-dimensional plane. When the cell attempts to leave the surface, it must deform the body into a sphere shape to detach the body from the surface and change the direction of the anterior tip, thereby liberating the flagellum, as shown in Fig. 3c, Fig. 3d, Fig. 1a, and Fig. 1b. When the cell comes near the contact line, the limited space restricts the rotation and beating of the flagellum (Fig. 3e). In the vicinity of a contact line, the wedge-shape of the space forces the cell move near the surface. The flagellum easily comes into contact with the substrate and is attached on the surface without active beating, in which case the flagellum malfunctions and cannot rotate (Fig. 3e). The motion of the cell stops, and it cannot turn around through changing the mode of rotation of the flagellum [8]. The head of the cell recedes, and its body exhibits metaboly, increasing the distance between the tip point and the contact line and pulling the flagellum backwards to recover its rotation, which spins the cell body. When the cell tip points backwards to a point where the cell can recover the beating and proper rotation of the flagellum, the cell escapes with flagellum-driven motion. Such an attempt does not always succeed in pulling the flagellum away from the attachment or restriction, and the cells that we observed typically carried out multiple attempts at deformation and rotation before they were successful. The complete malfunction of the flagellum also was found (Fig. 3f). Beating and rotation in the highly viscous condition hurt the flagellum. The flagellum is inactive and cannot rotate and beat. The cell undergoes metaboly continuously. The anterior tip contracts and draws back the flagellum, and the posterior part protrudes backwards. Such amoeboid behavior leads to a receding motion. The velocity is on the order of 1.0 μm/s. Additionally, when the flagellum breaks off, the cell immediately demonstrates the same receding movement. When the cell head is stuck on a substrate (Fig. 3g), the cell will deform continuously to try to pull back the head.

**Figure 3.**
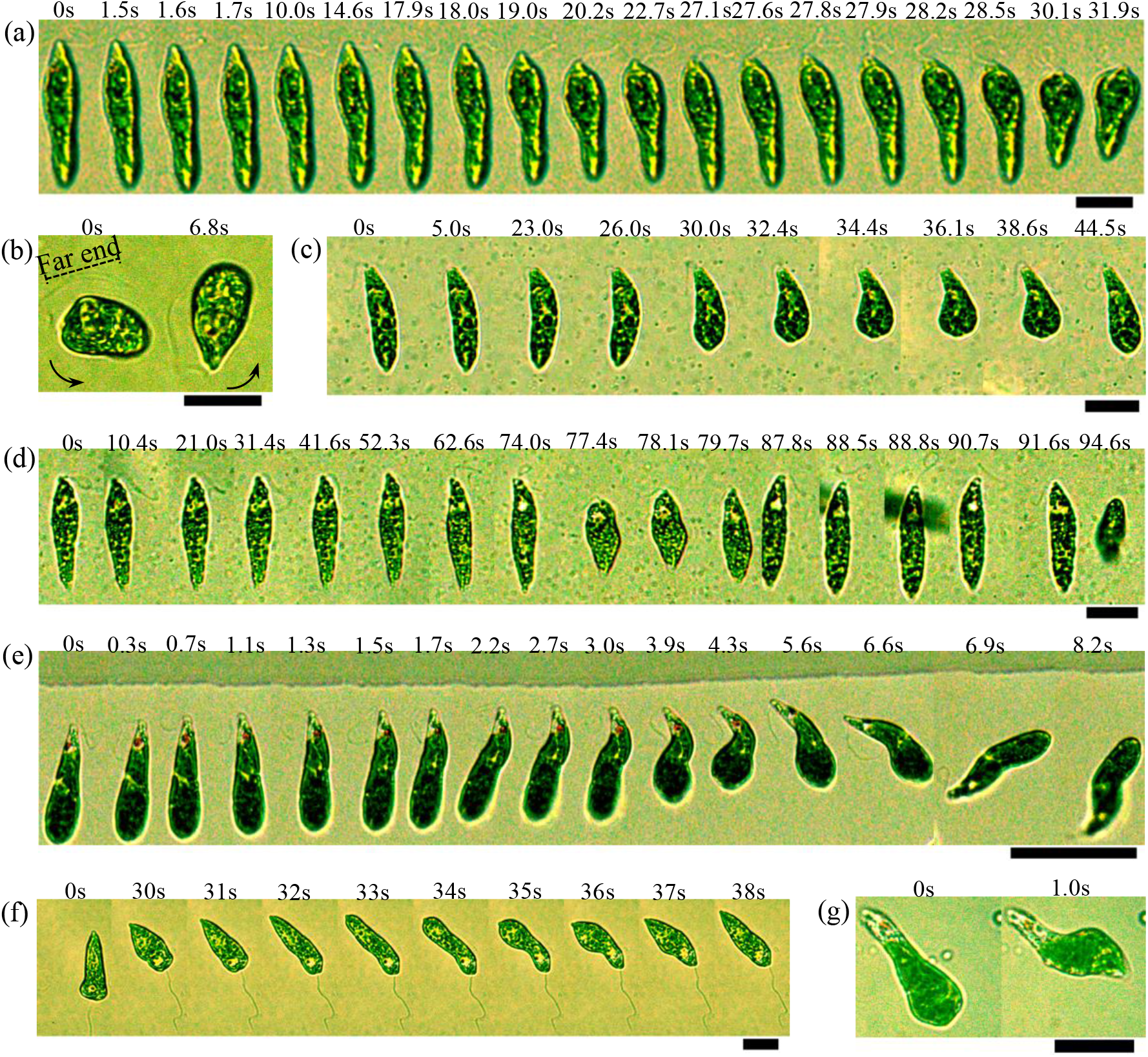
Flagellar malfunctions trigger metaboly. (a) Flagellum malfunction in a highly viscous solution (movie 14). The flagellum malfunctions from 0 s to 27.1 s. In (a), the concentration of the PAAS solution is 0.035wt%. (b) Rotations with metaboly when far end of the flagellum malfunctions (movie 15). The arrows indicate the directions of cell rotation. (c) Malfunction of the flagellum and metaboly when attached on a glass surface (movie 16). (d) Malfunction of the flagellum and metaboly when attached under the air-water interface (movie 17). The rotation of the flagellum is hindered by the water surface from 0 s to 74.0 s. (e) Malfunction of the flagellum, metaboly and escape in the vicinity of a contact line (movie 18). The flagellum malfunctions from 0.3 s to 6.6 s. (f) A cell with an inactive flagellum crawls backwards on a glass surface (movie 19). (g) Metaboly when the cell head is stuck to a substrate (movie 20). Scale bars: 20μm.

By examining the details of the flagellar motion, metaboly and escape process in the aforementioned conditions, we summarize that the flagellar malfunction triggers the metaboly in *E. gracilis*. The beating flagellum as a sensor detects threats in the surrounding environment. When the flagellum malfunctions, such mechanical information appears to be transduced into the cytoskeletal systems; the cell deforms its shape to decrease its body length to enlarge the space for the rotation and beating of the flagellum and turns round, driven by the rotation and beating of the flagellum. In this way, metaboly assists the cell in escaping from the restrictions. To reveal the functions of metaboly, we investigated advantages of metaboly under restrictions. When the cell is attached to a substrate or a wall and is in a cigar shape, the adhering force can be expressed as 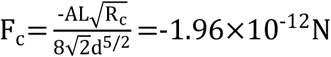, where the cell is seen as a cylinder adhering on a plane and A=3.7×10^−20^J, R_c_=4.5μm, L=50.0μm and d=0.5μm represent the Hamaker constant, radius of the cell, length of the cell and distance between cell and the surface [15] (Fig. 4a), respectively. After deformation into a sphere, the adhering force is 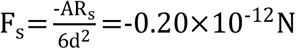, where the cell is viewed as a sphere adhering on a plane, and R_s_=8.2μm is the radius of the sphere (Fig. 4a) [15]. The ratio 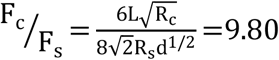 (Fig.4a), indicating that deformation from the cylindrical shape to a spherical shape substantially decreases the adhering force at the surface. For rotation (Fig. 4b), if the cell have cylindrical shape, the torque from the viscous resistance can be express as 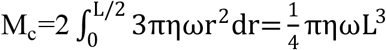, where *r* is the distance to the axis, η is the viscosity of water and ω is the angular velocity. If the cell deforms into a sphere (Fig. 4b), the torque from the viscous resistance can be approximately expressed as 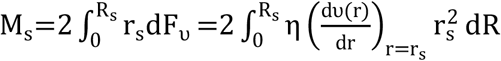, where F_υ_ is the viscous force on the surface, R is the distance to the center, υ(r) is the velocity of fluid, r is the distance to the axis and 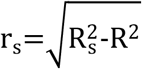. By solving the linear Navier-Stokes equation (Reynolds number is Re≤6.85×10^−5^), we have the velocity 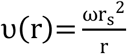 for a cylinder with length dR rotating in an infinite bulk fluid [16]. Thus, we have 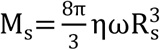. The ratio 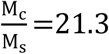, indicating that deformation from cylindrical shape to a spherical shape decreases the viscous torque substantially during rotation of the body. The cell can easily change the direction through metaboly in a harmful condition where the flagellum malfunctions. In addition, the deformation from cylindrical shape to spherical shape increases the radius of the cell and thus enlarges the space between the surface and flagellum (Fig. 4a(i)). In this way, the flagellum has a larger possibility of recovering its regular rotation and beating, which will produce enough torque and driving force to change the direction of the body and escape from the restriction of the adhering surface. However, if the deformation winds the cell tip towards the surface (Fig. 4a(ii)), it cannot recover the function of the flagellum. The cell will deform continuously to adjust its posture to that shown in Fig. 4a(i). Thus, metaboly plays key a role in restoring the function of flagellum and escaping from the restrictions.

**Figure 4.**
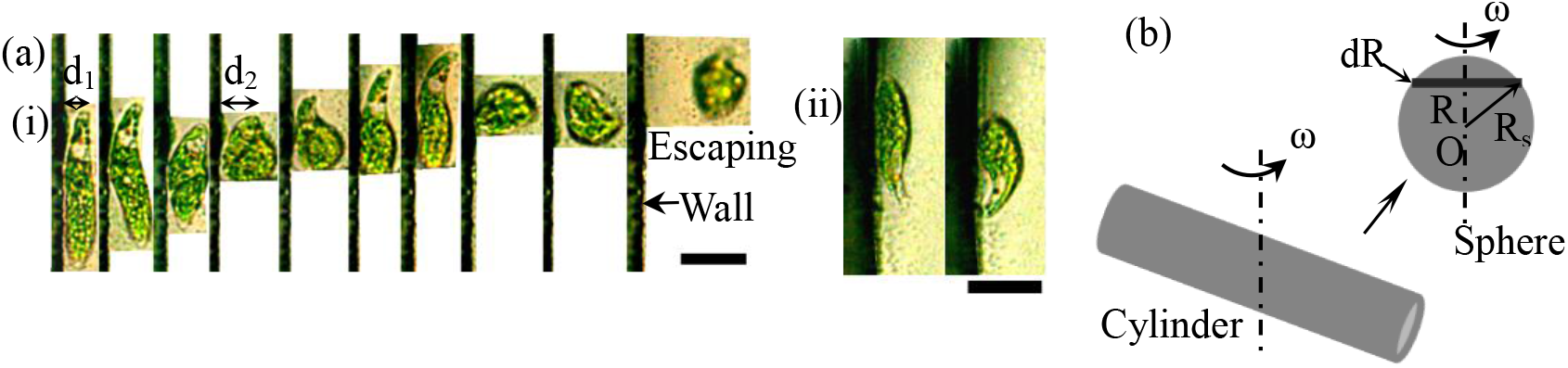
Functions of metaboly. (a) The cell attached on a wall and detachment. (i) detaching the body from the surface and decreasing the attaching area with the interface, which decreases the adhering force and is advantageous for escape (movie 21), (ii) winding the cell tip towards the wall surface (movie 22). The distances between flagellum and surface are d_1_=6.27μm and d_2_=11.08μm. Time interval is 0.3 s. Scale bars: 20μm. (b) Decreasing the torque of resistance for rotation of the body when deforming the body to from cylindrical shape to spherical shape.

Another question is how the cells take advantage of metaboly in the extreme case that the flagellum is completely out of function. Figure 5 demonstrates swimming in a bulk fluid and crawling on a surface. In the bulk fluid, the cell swims by metaboly when the flagellum breaks off as shown in Fig. 5a. In this case, the cell cannot rotate to change the direction of the body without the torque due to the rotation and beating of the flagellum. The cycles of retracting the head and then protruding the tail, over a period of approximately 7.0 s, produce net backward displacement in the fluid due to the nonreciprocal nature of the viscous force. Additionally, the cell can crawl fast on a surface through cycles of contracting the front part and protruding the rear part in the absence of confinement when the flagellum breaks off (Fig. 5b). The period of retraction and protrusion is also approximately 7.0 s. However, crawling on the surface is more effective than swimming in the bulk fluid, as swimming is susceptible to the effects of fluid flow and viscosity. In particular, when the body is spun by the beating of the flagellum (Fig. 2b), the cell rotates, and its trajectory is not a straight line. We also found that the cell crawls on a wet glass surface without bulk fluid (Fig. S5). On a wet surface, the flagellum cannot regularly rotate and beat as it does in the bulk fluid. The cell contracts at the front point, and then, the rear point protrudes. Such repeated deformation of the body produces net backward displacement on the surface with the assistance of the friction from the surface, with a velocity of approximately 1.5μm/s. The triggering mechanism is that the flagellum completely malfunctions or cannot rotate. Such crawling behaviour is useful for escaping (Fig. 5c&Fig. S3c): for example, when the head of a cell is stuck or penetrates something, this in turn inhibits rotation and beating of the flagellum and the cell pulls back its head successfully by crawling or swimming backwards (see Fig. 5c and movie 28). The cell crawls backwards to escape from the restrictions which are harmful for the rotation and beating of the flagellum. In addition, crawling behaviour in the confinement between two glass plates was observed, as shown in the Fig. 5d. When the live flagellum limited by two surfaces cannot rotate and almost completely malfunctions, the cell crawls backwards as it does when the flagellum completely malfunctions. Such metabolic crawling has previously been reported to be triggered by confinement [5]. Here, we show that confinement leads to malfunction of the flagellum, which then triggers metaboly, thus clarifying the mechanism underlying this earlier observation. However, when the flagellum can still rotate, the cell crawls forward, and the mode of deformation is inverse to that of backward crawling. The forward displacement is produced by the cycles of the head protruding and then the tail contracting. Usually, the forward crawling is combined with gliding.

**Figure 5.**
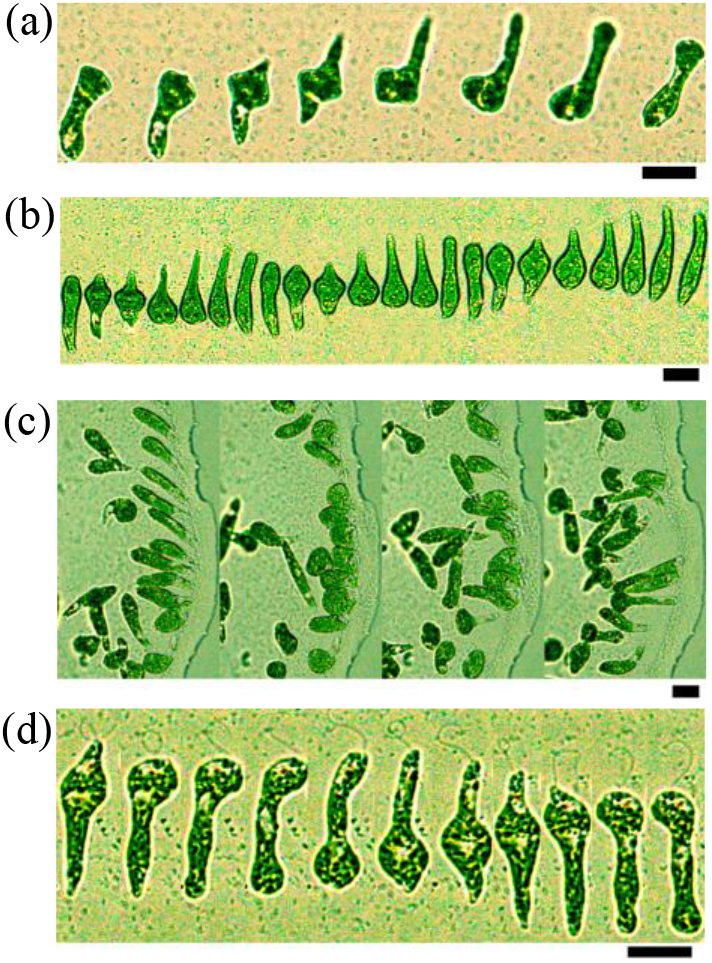
Metaboly-induced swimming and crawling. (a) Swimming backwards in a bulk fluid after the flagellum breaks off (movie 23). One period of metaboly is shown.(a) Crawling backwards on a glass surface when the flagellum completely malfunctions or break off (movie 24); (c) In a sticky situation and escaping (movie 25). The contact line is sticky due to a high concentration of the HUT culture medium because of evaporation of the solvent. The time interval is 9.0 s. (d) Crawling backwards when confined between two glass surfaces (movie 26). In (a), (b), and (d), the time interval is 1.0 s. Scale bars is 20μm.

In summary, we have solved the problems on the triggering mechanism and functions of metaboly in *E. gracilis*. *E. gracilis* cells demonstrate metaboly in response to flagellar malfunctions. Air-water interfaces, contact lines, walls, obstacles, high viscous conditions, and sticky environments are ubiquitous in the natural aquatic environments and restrict the regular functions of the flagellum of the cell. Without metaboly, the cell cannot escape from those restrictions. Thus, metaboly is an important escape strategy for the species. Our findings may provide insights on early euglenoid evolution. Furthermore, metabolic swimming and crowling without confinement in *E. gracilis* provide new hint for mechanism of metabolic motion [17–19]. Such a strategy can inspire the biomimetic design of soft self-adaptive robots replicating metaboly [20–22]. Additionally, it is possible to trap the cell physically using smooth surfaces with a large Hamaker constant, which produces a large adhering force toward the cells. Thus, we expect that our findings will have important implications for industry in the future.

## Methods

*Euglena gracilis* FACHB-227 was obtained from the Freshwater Algae Culture Collection at the Institute of Hydrobiology (FACHB, China). A 10ml *E. gracilis* solution was transferred into 200ml HUT culture medium (FACHB) in a flask that had been sterilized at 150 °C for twenty minutes, and the solution was cultured at a light:dark cycle of 12:12 hours under white illumination of two table lamps at room temperature (25°C) for a week. An upright microscope (Olympus BX53) with motorized stage was used for all experiments. The micrographs were recorded with a time-lapse CCD digital camera (Oplenic) fixed on the microscope at a frame rate from 10 f.p.s to 30 f.p.s. The data was analyzed using the image-analysis software ImageJ.

The glass plates and glass tubes used in the experiments were washed using detergent and water, then rinsed with ultrapure water (Millipore) and dried at 100°C. To observe the behaviour of cells at the air-water interface, the solution was deposited in an open chamber with a size of 25.0mm×5.0mm×0.57mm made by glass plates. A drop of solution containing the cell dispersion (~ 5μl) was deposited on a glass slide for observations. To observe the behavior of cell near a contact line, the microscope was focused on the contact area near the boundary of the droplet. The solution was transferred to a glass tube with a diameter of 7.14mm to observe the behaviour of cells attached to a glass wall and motion on a wet glass wall. To observe *E. gracilis* behaviour in extremely crowded conditions, a concentrated cell solution was collected from the vicinity of the air-aqueous surface of the culturing solution (the cells form a green film at the air-water surface). The aqueous solution of sodium polyacrylate (PAAS) (density: 1.32×10^3^ kg/m^3^, M_w_⩾3.0×10^7^, Sinopharm Chemical Reagent) was prepared by dissolving PAAS in Millipore water at 60°C. For observations of cells in high viscous conditions, cell solutions of high viscosity were prepared by gently mixing the HUT cell solution and PAAS solution at 25°C. The viscosity increases with the increase of the concentration of PAAS (Table S1). To observe the crawling behaviours of the cell in confinement, the cell suspension was drawn by capillary force into the two glass plates placed surface to surface.

## Acknowledgements

This work was supported by ZJNSF (No. LY16A040003) and NSFC (Nos. 11204181 and 11675112).

## Author contributions

Y.-J. C. designed the research, performed the experiments, analysed the data and wrote the manuscript.

## Materials and Correspondence

should be addressed to Y.-J. C.

## Extended Data

**Extended Data Figure S1.**
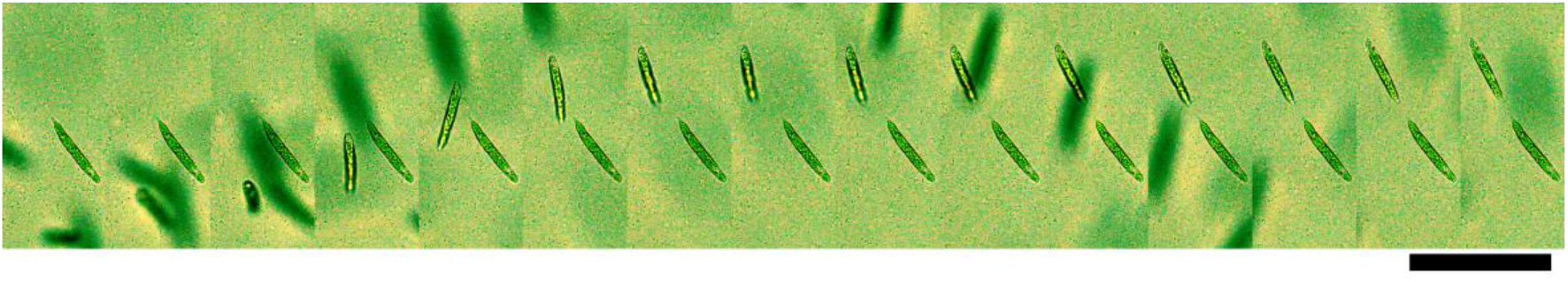
The air-water surface traps the cell (movie 5). The time interval is 0.5 s. Scale bar: 100μm.

**Extended Data Figure S2.**
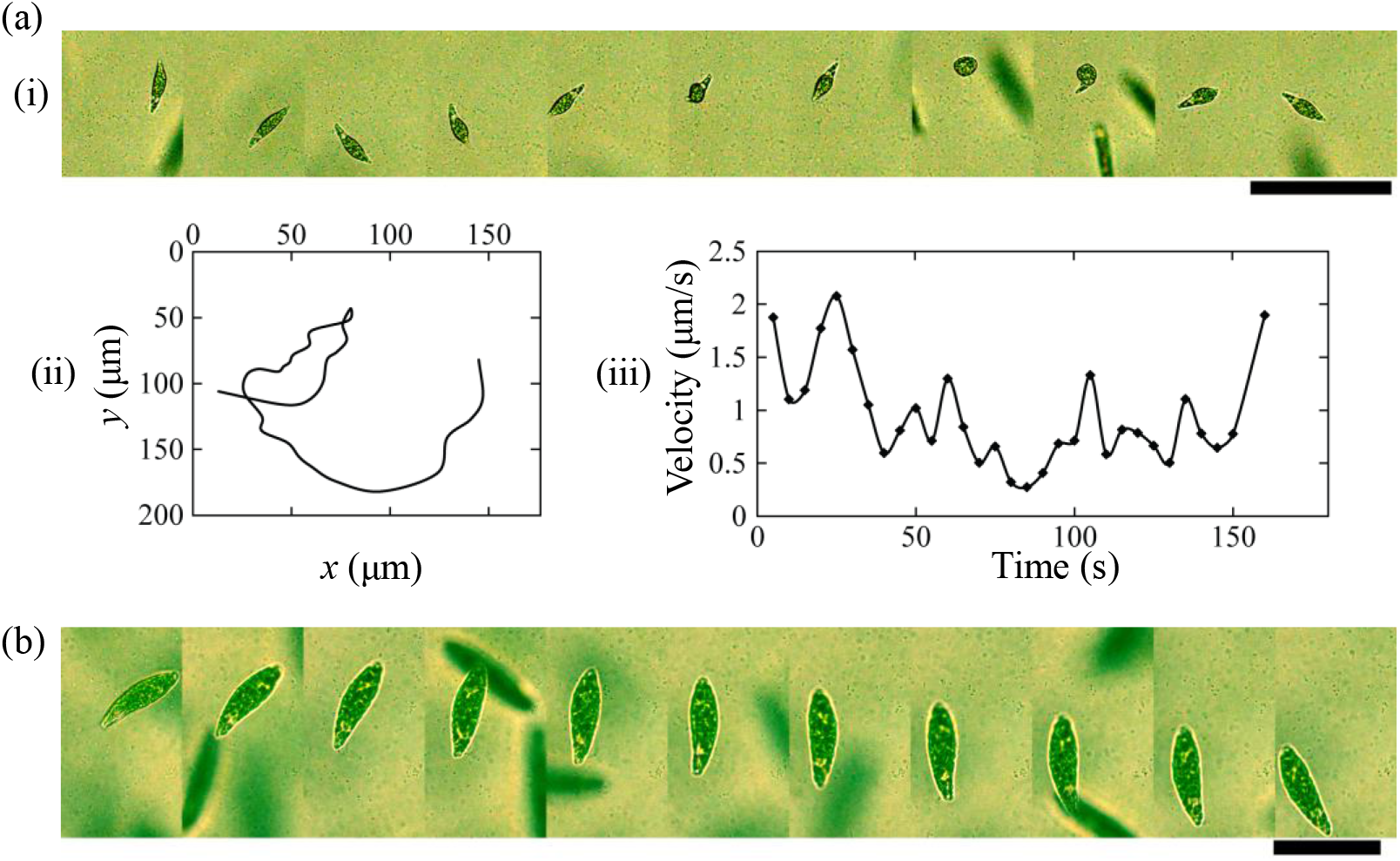
Gliding on a surface or along the air-water interface. (a) The attached cell glides under the air-water surface (movie 6). (i) Spatial-temporal displacement of cell gliding under the air-water surface, with a time interval is 15.0 s. Scale bar: 100μm. (ii) Trajectory of the cell gliding. (iii) Time-dependent velocity of cell gliding under the air-water surface. (b) The cell glides on a glass substrate (movie 7). The time interval is 2.0 s. Scale bar: 50μm.

**Extended Data Figure S3.**
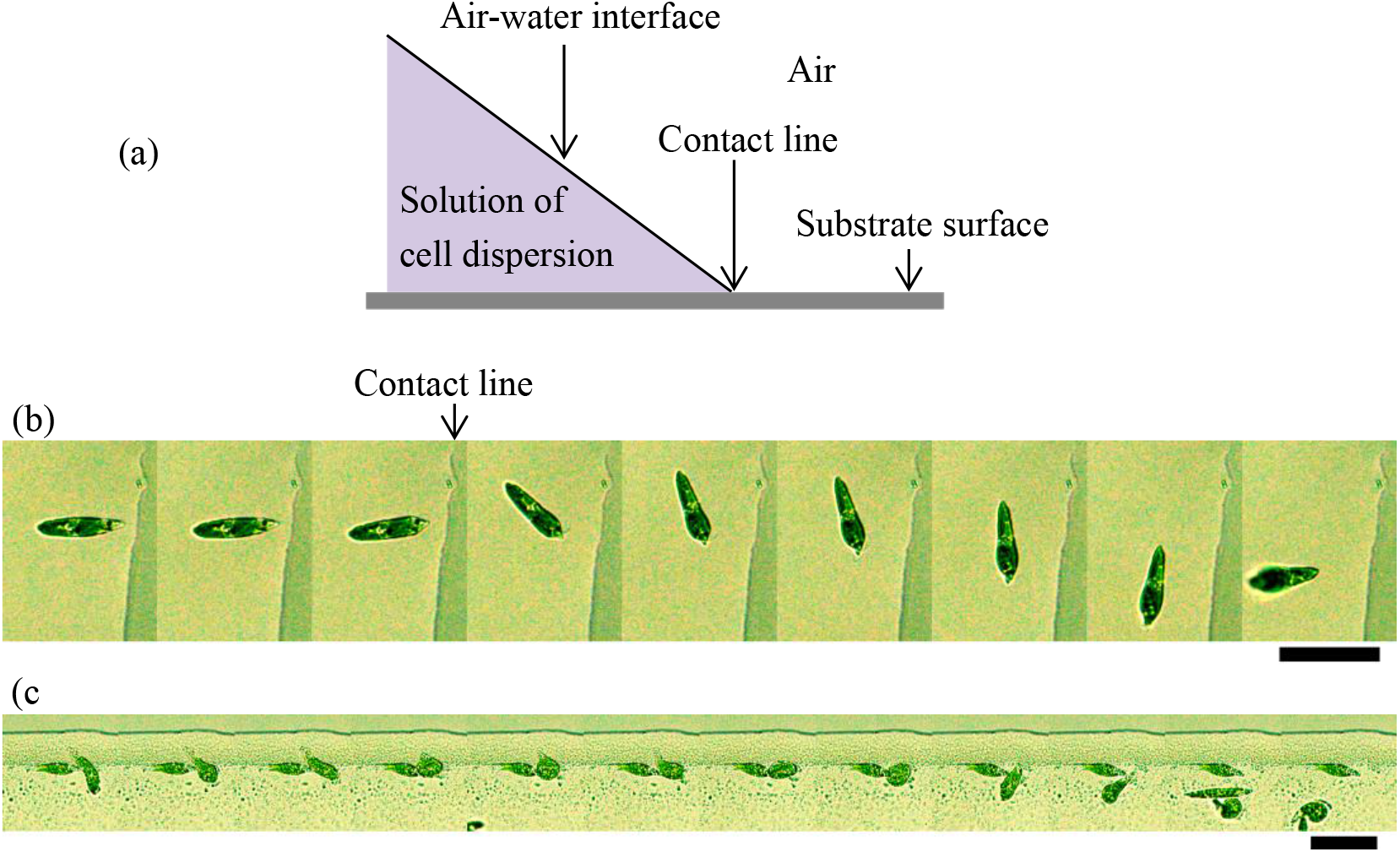
A cell uses metaboly to turn around. (a) Schematic diagram of contact line. (b) The cell turns around and escapes through slight deformation (movie 28). The time interval is 0.5 s. (c) Metaboly when the anterior part of the cell is stuck, and subsequent escape (movie 29). The time interval is 2.0 s. Scale bars: 50μm.

**Extended Data Figure S4.**
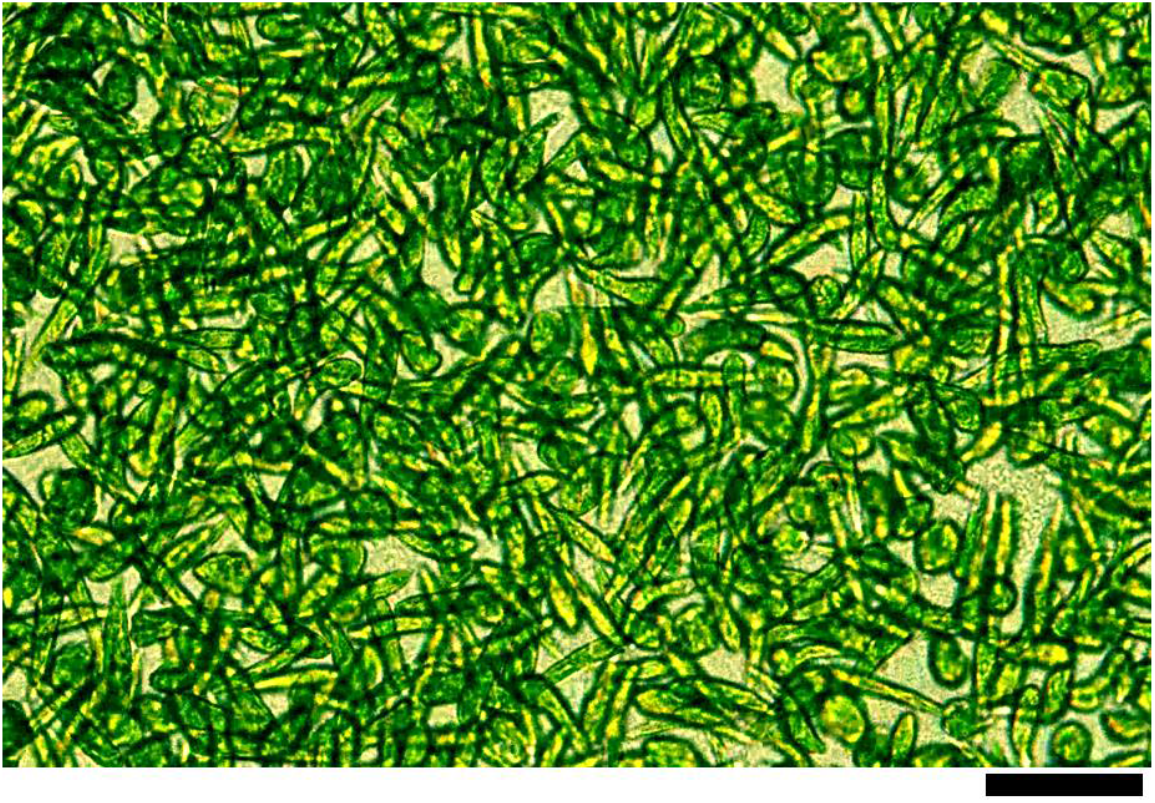
Metaboly in an extremely crowded environment (see movie 9). Scale bar: 50μm.

**Extended Data Figure S5.**
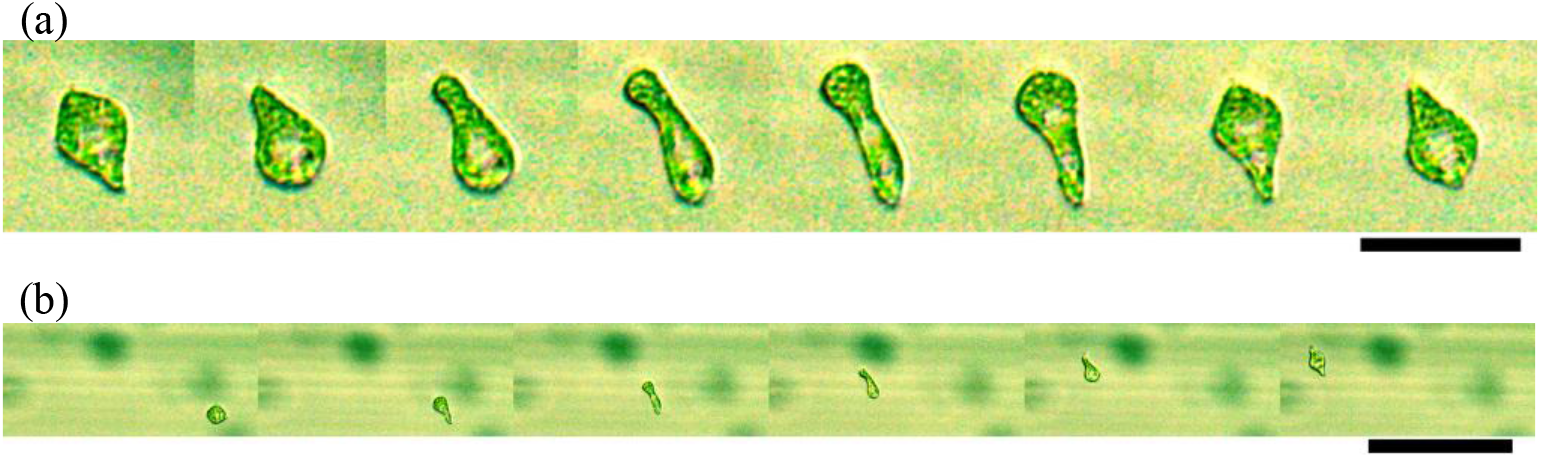
Crawling on a wet glass surface (movie 27). (a) One period of metaboly during crawling. The time interval is 1.0 s. Scale bar: 50μm. (b) Positions of cell during crawling on the wet surface. The time interval is 5.0 s. Scale bar: 200μm.

**Extended Data Table S1.** Measurement of viscosity of diluted PAAS aqueous solution at 25°C. The viscosity of PAAS aqueous solution was measured using a digital rotary viscometer (NDJ-5S, Lichen Science and Technology, China). According to the measurements, the PAAS aqueous solution is a non-newtonian fluid.

| Concentration (wt%) | Number of rotator | Rotation rate (Rotation Per Minute) | Viscosity (mPa·s) |
| --- | --- | --- | --- |
| 0.1000 | 1 | 6 | 578.0 |
|  |  | 12 | 341.5 |
| 0.0700 | 1 | 6 | 390.9 |
|  |  | 12 | 232.0 |
| 0.0500 | 1 | 6 | 257.0 |
|  |  | 12 | 156.5 |
|  |  | 30 | 84.00 |
|  |  | 60 | 70.00 |
| 0.0350 | 1 | 6 | 181.0 |
|  |  | 12 | 113.5 |
|  |  | 30 | 62.60 |
|  |  | 60 | 42.90 |
| 0.0250 | 1 | 6 | 135.0 |
|  |  | 12 | 84.50 |
|  |  | 30 | 47.80 |
|  |  | 60 | 33.50 |
| 0.0250 | 0 | 6 | 45.20 |
|  |  | 12 | 29.15 |
|  |  | 30 | 17.40 |
| 0.0170 | 0 | 6 | 29.90 |
|  |  | 12 | 19.90 |
|  |  | 30 | 12.10 |
|  |  | 60 | 8.80 |
| 0.0100 | 0 | 6 | 17.90 |
|  |  | 12 | 12.20 |
|  |  | 30 | 7.64 |
|  |  | 60 | 5.78 |
| 0.0050 | 0 | 6 | 11.2 |
|  |  | 12 | 7.44 |
|  |  | 30 | 4.52 |
|  |  | 60 | 3.55 |
| 0.0025 | 0 | 30 | 2.88 |
|  |  | 60 | 2.37 |
| 0 | 0 | 60 | 1.05 |

